# Film Recall Reveals Intact Event Memory but Impaired Sequence Memory in Temporal Lobe Epilepsy Patients

**DOI:** 10.64898/2026.02.03.702007

**Authors:** Herui Zhang, Forouzan Farahani, Eden Tefera, Zayn Ahmed, Benjamin Botnick, Bijay Thapaliya, Hongmi Lee, Helen Borges, Wenze Zhang, William Barr, Simon Henin, Yidan Shi, Janice Chen, Anli Liu

**Author notes:** Corresponding Author: Anli Liu MD MA. Shared first-authorship. Shared senior authorship. Disclosures: None.

## Abstract

**BACKGROUND AND OBJECTIVES:** Patients with epilepsy (PWE), especially temporal lobe epilepsy (TLE), experience impaired memory for personally experienced events. However, current assessments of episodic memory are limited in their ecological validity with a potential to miss detection of subtle cognitive decline. We conducted an exploratory study to determine whether a naturalistic film-viewing task with open-ended spoken recall could detect memory differences between TLE patients and healthy controls (HCs).

**METHODS:** TLE patients (ages 18-60, fluent in English, not legally blind) were recruited from a Level 4 Epilepsy Center (2018-2024). TLE diagnosis was based on seizure semiology, MRI Brain, and EEG. TLE patients scored ≥22/30 on the Montreal Cognitive Assessment (MOCA); HCs scored ≥26/30. Subjects watched 6 short films and then freely recalled film details. Spoken responses were recorded, transcribed, segmented, and scored for film- and event-level recall. Recall order was assessed using the Damerau-Levenshtein distance. Semantic and causal centrality were quantified using sentence embeddings and rater-identified causal links, respectively. Beta regression with cluster-robust standard errors assessed group and centrality effects on recall probability. Beta regression evaluated the influence of age, MOCA, and testing platform on sequence recall error.

**RESULTS:** We recruited 51 subjects (27 TLEs; 24 HCs, 70.1% F, mean 29.9 ±8.3 years). TLE patients and HCs showed similar recall of films (HC 89% ±11% vs TLE 88% ±18%, p = 0.54), coarse events (HC 50% ±16% vs TLE 44% ±18%, p = 0.19) and fine events (HC 25%±10% vs. TLE 22%±12%, p=0.17). Both groups recalled high causal centrality events better. For coarse event sequence recall, TLE patients showed a numerical trend toward greater sequence errors compared to HCs (HC **10.8% ± 10.5%** vs. TLE **19.5% ± 18%**, *p* = **0.06**), although this difference did not reach statistical significance. However, TLE patients showed significantly greater fine event sequence errors at recall than HCs (HC 15% ±13% vs 23% ±18%, p = 0.02, Hedges’ *g* = 0.85, Cliff’s δ = 0.51), with RTLE demonstrating more sequence errors than HCs (15%±13 vs. 29%±21% p = 0.021) Age, education, MOCA, and performance on standard verbal and visual memory tasks were unrelated to film, event, and sequence recall performance.

**DISCUSSION:** We demonstrate that a short film task with spontaneous spoken recall can identify sequence memory impairment in TLE patients despite intact film- and event-level recall. Sequence memory may represent a subtle manifestation of memory impairment that is not detected by standard cognitive testing.

**Key Points:**

- We asked whether a naturalistic film recall task could detect episodic memory impairment in a temporal lobe epilepsy cohort.
- Patients with temporal lobe epilepsy showed comparable film and event recall compared to healthy controls but were found to have impaired sequence memory.
- Sequential memory for temporal order is an overlooked aspect of episodic memory that may detect subtle memory decline.

## INTRODUCTION

Episodic memory, or the ability to remember past events, plays a critical role in our ability to remain oriented to the present and plan for the future (1). Memory is often impaired in neurological disorders like epilepsy, traumatic brain injury, and Alzheimer’s Disease, impacting patients’ ability to work and live independently. One major limitation in translating findings from basic memory research is that test behavior is studied in an artificial manner. For example, rote or unstructured verbal memory is commonly assessed using arbitrary lists of items (e.g., words, objects, faces) without context, while assessment of structured verbal memory is mostly limited to story recall (2–4). This traditional approach to memory testing has provided valuable contributions but overlooks important aspects of real-world memory, cognition, and behavior, including visual, auditory, and sequence information (5). Naturalistic cognitive testing holds potential for assessing episodic memory dysfunction with greater ecological validity in clinical populations and potentially revealing subtle aspects of memory impairment (6).

Films have been used in cognitive neuroscience to examine the neural dynamics underlying naturalistic perception and memory. Because films present a continuous multi-modal stream of sensory information, they are thought to elicit cognitive processes resembling those of daily life (7). The brain must segment a continuous flow of visual, auditory, and tactile information into meaningful units, a process known as event segmentation (8,9). This automatic process of chunking, storing, and retrieving information occurs similarly across humans at both behavioral and neural levels; healthy subjects spontaneously recall film events in the order of presentation and denote similar event transitions (10–14). Hippocampal activation and other default mode network (DMN), such as posterior cingulate, precuneus, parietal regions, are observed at event boundaries (15,16). Events with high semantic centrality and causal centrality—those that are interconnected with many other events—are more likely to be remembered (17–19). Because films mimic the continuous, multimodal nature of everyday memory function, their use in cognitive testing may enable the detection of key features of episodic memory.

Building on our prior findings, we aimed to determine whether naturalistic film-viewing could be used to identify episodic memory impairments in TLE patients. It is now acknowledged that TLE patients demonstrate a range of cognitive phenotypes. Some have normal cognition, others have memory and/or language impairment, and some have global or multi-domain impairment (20,21). Approximately 70% of TLE patients report memory dysfunction (22), but only half demonstrate objective deficits after the standard delay of 30 minutes during traditional clinical testing (23). Accelerated forgetting occurs after normal encoding and after typical 30 minute test delays, when TLE patients are retested at 1 to 3 week intervals (24–26). Accelerated forgetting is thought to be due to abnormally fast decay of initially remembered information and disrupted sleep-dependent consolidation (27).

We focused on our task’s ability to detect subtle differences at the immediate recall stage. By presenting short films to both TLE patients and healthy controls, followed by free recall, we examined differences in naturalistic film-level, event-level, and event-sequence recall. We hypothesized that TLE patients would demonstrate poorer film, event, and sequence-level recall compared to HCs.

## MATERIALS & METHODS

### Subjects

Patients were recruited from a Level 4 Epilepsy Center in New York City (2018-2024). Eligibility criteria included a diagnosis of probable or definite unilateral TLE if they possessed concordant (1) seizure semiology, (2) MRI Brain abnormalities, and/or (3) ictal or interictal EEG, as confirmed by a board-certified epileptologist (AL). TLE patients were eligible if they scored ≥22 out of 30 on the Montreal Cognitive Assessment (MoCA), a brief screening questionnaire that assesses attention, memory, naming, and visuospatial functioning. HCs were recruited from the community, eligible if their MOCA score ≥ 26 and did not have a history of neurological or psychiatric disorders. We selected a lower MOCA threshold for TLE patients to capture a greater range of cognitive and memory performance. In our analysis, we used the total MOCA score (MOCA Total) and the score from the MOCA delayed recall trial (MOCA Delay).

In a secondary analysis, we compared film recall to standard neuropsychological testing for the subset of TLE patients who had available testing performed for clinical purposes. Between 30-60% of TLE patients had test results available for the Logical Memory and Visual Reproduction subtests from the Wechsler Memory Scale (WMS-IV) (28), Rey Auditory Verbal Learning Test (RAVLT) (29), and Rey Complex Figure Test (RCFT) (30), depending on the task. Variables of interest included the immediate and delayed trials of the WMS-IV Logical Memory (LMI & LMII) and Visual Reproduction (VRI & VRII) subtests. RAVLT variables included total learning over five trials (RAVLT Total), short delayed recall (RAVLT VI), and long delayed recall (RAVLT VII). Analysis of the RCFT was limited to the delayed recall trial (RCFT DR) and the analysis of the MAC-E was limited to the total frequency score (MAC-E Total) (31).

### Protocol and Consent

All study procedures were approved by the NYU Institutional Review Board, and written informed consent was obtained from every participant.

### Naturalistic Film Task

The task consisted of a series of five short audiovisual films, including one animation and four live-action films, similar to our previously published paper (18)(**Figure 1A**). The films presented diverse narrative content and structures and ranged from 2.23 to 5.82 minutes in duration (avg. 4.03 min). Each film began with a 6-second image of the title in white letters on a black background. Each film was displayed on a separate white screen and manually advanced to the next with a mouse click. The order of the films was the same for all participants. A short cartoon (“Intermission”, 38 seconds) was shown at the beginning of each viewing session but was excluded from the analysis. Films are described in further detail in **Supplementary Table 1**.

**Table 1.** Subject Demographics, Clinical Features, and Film Recall Performance.

| Diagnosis | Healthy (24) | TLE (27) | Total (51) | p value |
| --- | --- | --- | --- | --- |
| **Age, years (SD)** | 27.6 (7.9) | 32.0 (8.1) | 29.9 (8.3) | 0.002* |
| **Sex** |  |  |  |  |
| Female | 18 | 18 | 36 | 0.73 |
| Male | 6 | 9 | 15 |  |
| **Handedness** |  |  |  |  |
| Right | 22 | 21 | 43 | 0.29 |
| Left | 1 | 3 | 4 |  |
| Ambidextrous | 0 | 2 | 2 |  |
| **Education Level** |  |  |  |  |
| 13-15 years | 2 | 4 | 6 | 0.7 |
| 16 years | 11 | 10 | 21 |  |
| ≥17 years | 11 | 13 | 24 |  |
| **MOCA (out of 30, SD)** | 28.6 (1.1) | 26.6 (1.8) | 27.6 (1.8) | <0.001*** |
| **MOCA Delay Recall (out of 5, SD)** | 4.66 (0.6) | 3.653 (1.2) | 4.14 (1.12) | <0.001*** |
| **Film Recall Probability % (SD)** | 89% (11.7%) | 88% (18.6%) | 89% (15.6%) | 0.54 |
| **Coarse Event Recall Probability % (SD)** | 50.4% (16.7%) | 44.6% (18%) | 47.3% (17.5%) | 0.19 |
| **Fine Event Recall Probability % (SD)** | 25.2% (10%) | 22% (12.3%) | 23.5% (11.3%) | 0.17 |
| **Film Sequence Recall Error % (SD)** | 15.8% (27.2%) | 23% (37.7%) | 20% (33.1%) | 0.42 |
| **Coarse Event Sequence Recall Error % (SD)** | 10.8% (10.5%) | 19.5% (18%) | 15.4% (15.4%) | 0.060 |
| **Fine Event Sequence Recall Error % (SD)** | 15% (13.7%) | 23.8% (18.1%) | 19.7% (16.6%) | 0.027 |

**Figure 1.**
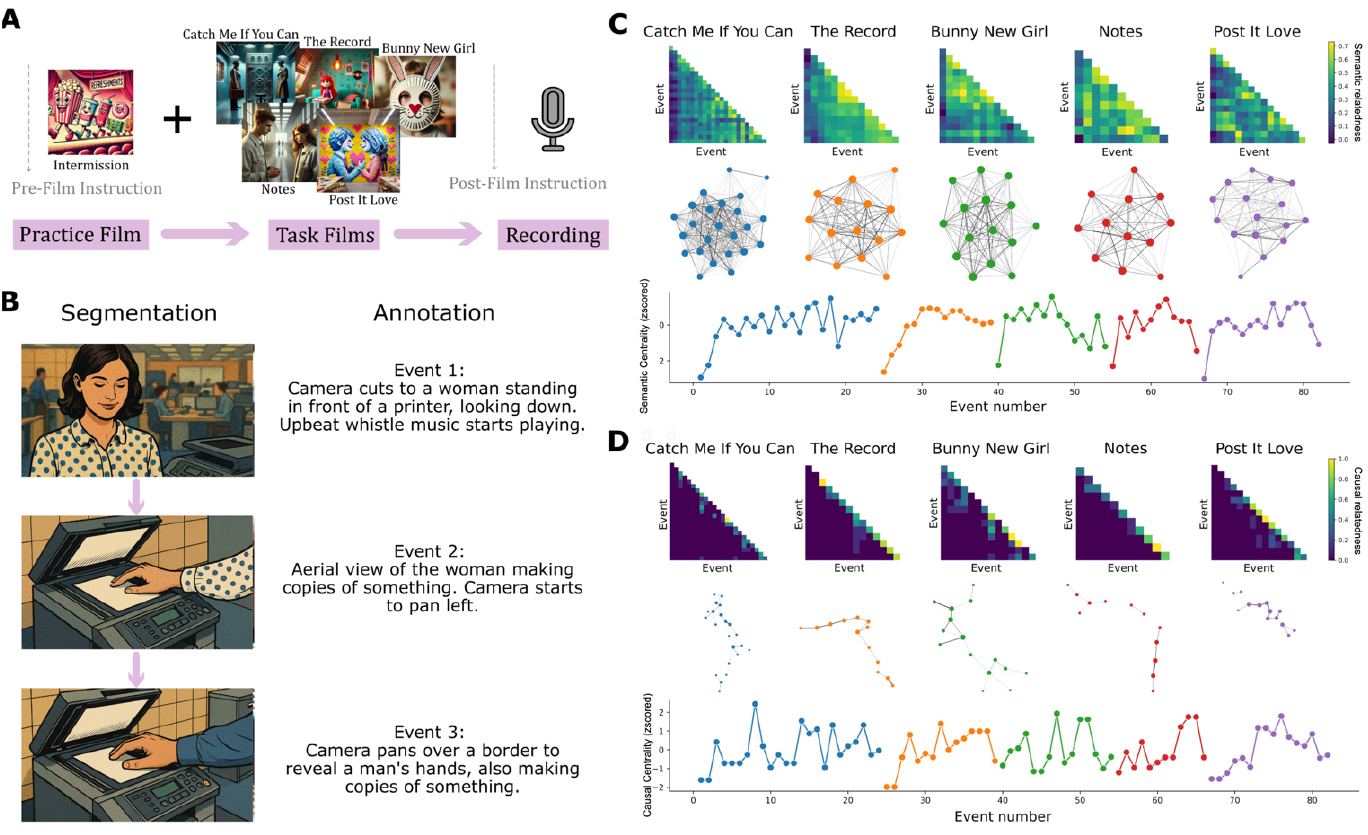
Naturalistic Film Task. A. Task design. Subjects were shown an introductory “practice” film followed by five “task” films. After viewing all films, subjects were instructed to recall aloud as much as they could remember about each of the films. Due to copyright constraints, all images in this figure were generated by ChatGPT and are not actual screenshots from the film. B. Event segmentation and recall scoring. Each film was segmented into events, with event boundaries determined by shifts in the narrative. Subjects’ spoken responses were recorded, transcribed, and scored for whether they remembered each film’s events, and sequence of events. C. The semantic centrality matrix (top), semantic narrative network (middle), and z-scored event time course (bottom) for each film. D. The causal centrality matrix (top), causal narrative network (middle), and z-scored event time course (bottom) for each film.

Subjects participated in one of two environments: (1) in person at the NYU Comprehensive Epilepsy Center, or (2) remotely through WebEx, a HIPAA-compliant video conferencing application. WebEx was introduced as a testing platform in 2020 during the COVID-19 pandemic, during which all research activities were conducted remotely. The task consisted of two phases: film watching and open-ended recall **(Figure 1A)**. Subjects were instructed to watch the series of short films carefully and told that they would later be asked to recall the films^18^. Subjects were then encouraged to freely describe everything they remembered about the films and to speak for at least 5 minutes.

### Event Segmentation & Annotation

Each of the five films was segmented into 12-24 coarse events (mean: 16.4 events) **(Supplementary Table 1)**. Coarse event boundaries were identified by an independent rater based on significant narrative shifts such as changes in location, topic, or time (**Figure 1B**). This method aligns with established protocols where coders mark transitions in the narrative to delineate event boundaries (10,18). There was no minimum duration required for an event; events lasted between 1 and 42 seconds, with an average of 14.9 seconds. Three independent raters provided detailed annotations of each event. In addition to these coarse boundaries, the same independent annotators identified fine event boundaries. Fine-grained, or detailed, event segmentation was performed by marking boundaries within each coarse event based on their own judgements of moment-to-moment narrative shift. Across the five films, the number of fine events ranged from 34 to 59, with a mean of 45.4 events per film (SD=10.38). Raters annotated what was happening within each fine event. The beginning and end of each fine event were timestamped within the scoring rubric. Example coarse and fine event annotations are available in **Supplementary Table 2**. Coarse and fine events allow for different analyses: while we go on to assess memory and semantic relations between events at both scales, causal labeling was performed at the coarse scale only. This is because causal labeling must be performed via human judgments, and the number of event-pair comparisons required for the fine event scale would be prohibitive. E.g, for the movie Catch Me If You Can, there were v=24 coarse events and v=59 fine events; the number of event-pair comparisons is (v*(v-1))/2; 276 comparisons are needed for coarse events, 1711 are needed for fine events.

### Recall Transcription

Subjects’ audio recordings were transcribed using automated and manual methods. Transcripts were first generated through Whisper AI (for in-person testing) or WebEx (for remote testing). These automated transcripts were verified and corrected by a member of the research team (HZ).

### Recall Scoring

Recall scoring was conducted manually by two independent scorers (HZ, ZA), who matched event annotations with subject-generated event descriptions seen in recall transcripts (**Figure 1B**). Consistency was ensured through adherence to a predefined scoring protocol. For the initial five subjects, both scorers independently evaluated the responses and subsequently reconciled their scoring criteria through discussion.

To determine the timing of these segments during recall, we used the *whisper_timestamped* package (based on Whisper AI) (32–34) to obtain word-level timestamps for each event segment’s start/end. We then applied the *SequenceMatcher* class from Python’s *difflib* module to align the recall word sequences with the corresponding events, enabling us to extract the precise timing for each segment.

Film recall probability was calculated by dividing the number of films recalled by the total number of films (5). For both coarse and fine events, recall probability was determined by dividing the number of events recalled by the total number of events in each film. We then averaged this ratio across all five films to weigh short and long films equally. Memory for film and event sequences was assessed using the Damerau-Levenshtein distance, which calculates the minimum number of edits (insertions, deletions, substitutions, or transpositions of adjacent items) required to match one sequence to another (35). Because it treats transpositions of adjacent items as a single edit, it is particularly sensitive to subtle positional errors in recall tasks, where swapping two neighboring items should be penalized less severely than multiple separate edits. For film-level recall order analysis, we compared the recalled movie sequence to the actual film sequence (1, 2, 3, 4, 5). A similar method was applied for event-level recall order. One challenge in grading sequence performance is that poorer event memory may be erroneously associated with better sequence memory because there are fewer opportunities for errors. To correct for this, the Damerau-Levenshtein distance is normalized by dividing it by the total number of items recalled, generating a normalized deviation score for sequence memory that is independent of the number of films or events remembered.

### Semantic and Causal Centrality Scoring

An event’s semantic centrality reflects how strongly it is connected overall to other events in the network, e.g., events with high semantic centrality are strongly connected to (share features with) multiple events in the network. For example, two events that take place in a library (shared feature of location) will have higher semantic similarity than if one of the events takes place in a library and the other event takes place in a restaurant. To calculate the semantic centrality of film events, we converted the film annotations to sentence embeddings using Google’s Universal Sentence Encoder (USE) (36), as in our prior work **(Figure 1D)**. USE generates 512-dimensional embeddings that capture contextual meaning, enabling semantic comparisons of narrative content (36). Each film event was transformed into a 512-dimensional vector. A narrative network for each film was constructed by calculating the cosine similarity between pairs of these event vectors, which served as edge weights in the network, representing the semantic relationships between events (18). Semantic centrality was calculated as the sum of all of an event’s edge weights. These analyses were performed separately for the fine events and the coarse events.

An event’s causal centrality reflects the overall strength of its causal connections to other events in the narrative. A scene in which the protagonist steals a key that later permits a break-in, triggers an alarm, a chase, and an arrest possesses high causal centrality, whereas a brief moment where a character pauses to retie a shoelace without consequences is low in causal centrality. For causal centrality analysis, 13 independent raters were tasked with identifying causal relationships between coarse event pairs within each film. Raters viewed the entire film and were provided with detailed annotations of the events. Two events were labeled as being causally connected if what happened in one could explain what happened in the other, regardless of the order of occurrence; this was recorded as an edge weight of 1 (causally connected) or 0 (not causally connected). An event’s causal centrality was calculated as the sum of all edge weights per event, averaged across raters. An event’s causal centrality reflects the overall strength of its causal connections to other events in the narrative. Detailed rater instructions are provided in the **Supplementary Methods**.

### Statistics

We performed descriptive statistics for categorical variables such as sex, handedness, and education level. A chi-squared test was applied to the categorical data for comparison. For continuous data such as age, MOCA scores, film and event recall percentage, and sequence error, we used the Shapiro-Wilk test to test for normality. For normally-distributed data with comparable variances between the HC and TLE groups, such as the event recall probability, we employed the Two-Sample T-Test. For data that were not normally distributed, such as movie recall probability, we utilized the Mann-Whitney U Test. Additionally, we conducted post-hoc analysis using the Kruskal-Wallis Test for non-parametric comparisons between HCs and TLE patients, then for post-hoc analysis between HC, LTLE, and RTLE groups. Recall probability was analyzed with beta regression including centrality (High vs. Low), group (Healthy vs. TLE), and their interaction as predictors. Cluster-robust standard errors by subjects accounted for repeated measures, and Wald χ^2^ tests were used to evaluate main effects and the interaction. Beta regression was used to address potential mediators or confounds for outcomes, such as the contribution of age, MOCA score, TLE diagnosis, and education level on sequence memory performance. Effect sizes are reported for normally distributed data with Hedges’ g and for non-normally distributed data as Cliff’s δ.

## RESULTS

A total of 51 subjects (27 TLE, 25 HCs; 15 RTLE, 12 LTLE) were recruited. Thirty patients (58.8%) tested in-person, and 21 (41.2%) tested remotely. Groups were matched by sex (p = 0.731), handedness (p =0.292), and education (p = 0.702) (**Table 1**). Significant differences were observed between HC and TLE on age (27.6±7.9 vs 32.0 ± 8.1 years, p=0.002) and MOCA testing (28.6 ±1.1 vs 26.6 ±1.8, p = 0.00004), with the latter group difference due to different eligibility criteria (Table 1).

### TLE Patients Demonstrate Similar Film and Event Recall Performance Compared to HCs, but Poorer Sequence Recall

We anticipated that TLE patients would exhibit poorer memory performance, manifested by a lower recall probability and a higher sequence recall error. There were no significant differences at the group level between TLE patients and HCs in film recall probability (HC 0.89 ±0.11 vs TLE 0.88 ±0.18, p = 0.54) (**Figure 2C)**. TLE patients did not differ significantly from HCs for memory of coarse events (HC 0.50±0.16 vs TLE 0.44±18, p =0.19) or fine events (HC 0.25±10 vs TLE 0.22±0.12, p=0.17) (**Table 1, Figure 2D**). Inspection of the raw data of coarse and fine event recall performance between all patients reveals wider heterogeneity among TLE patients compared to HCs (**Supplementary Figure 3)**. We did not find a difference between recall for events during the first or last films for either HCs or TLE patients (p=0.629, see **Supplementary Figure 5)**. Likewise, there was no significant difference in the order of films recalled between the two groups (HC 0.15 ±0.27 vs TLE 0.23 ±0.37, p = 0.42) (**Table 1, Figure 2C**). Both coarse and fine event-level recall were comparable between groups **(Table 1, Figure 2D)**. TLE patients demonstrated significantly more sequence errors for fine events compared to HCs (HC 0.15±0.13 vs TLE 0.23±0.18, p=0.027); TLE patients showed a trending difference in sequence memory for coarse events compared to HCs (HC 0.10±10 vs TLE 0.19±0.18, p=0.060) (**Table 1, Figure 2E**).

**Figure 2.**
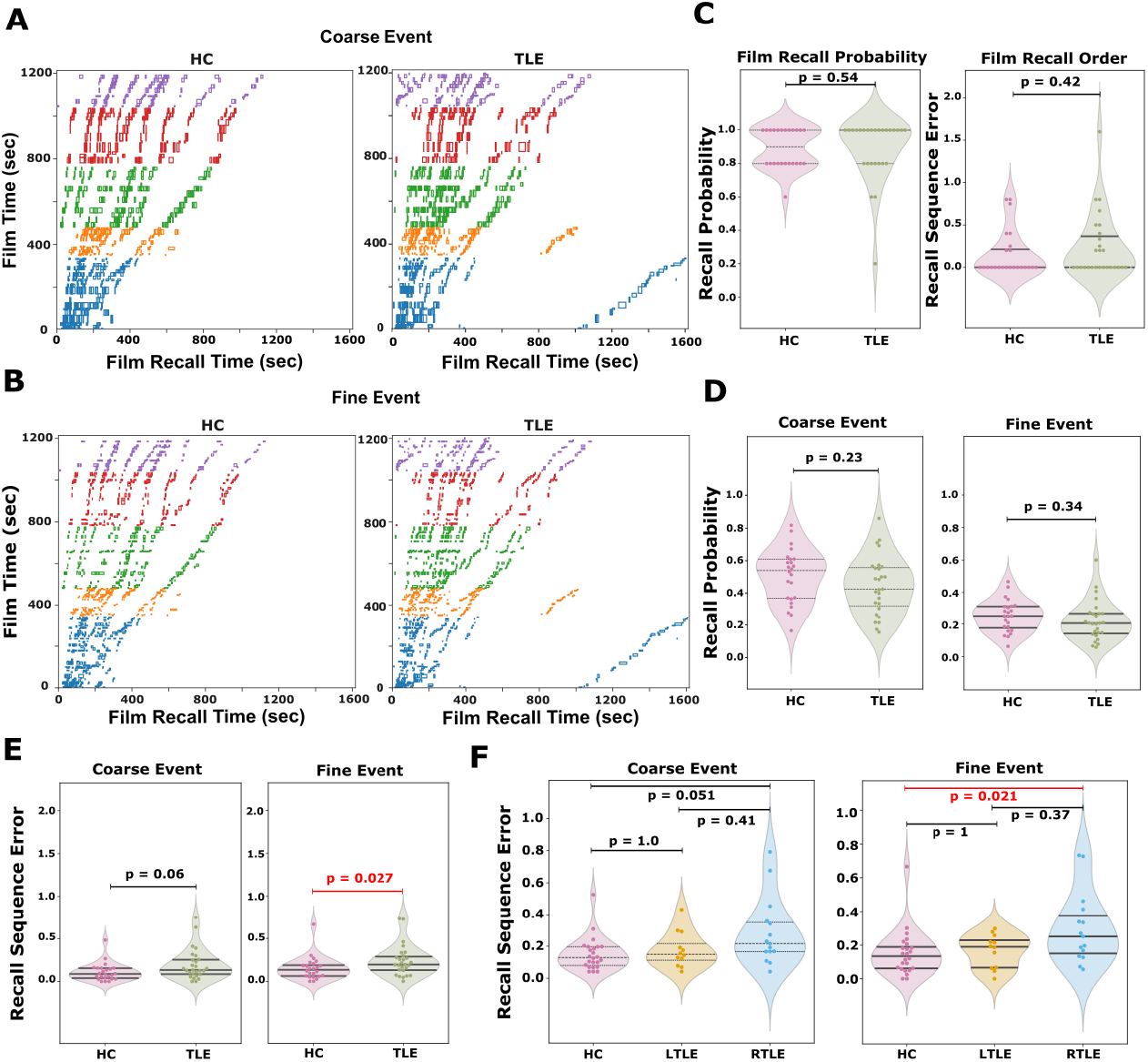
Recall performance and visualization. **A. Temporal correspondence between spoken recall and presentation order of coarse film events.** Left panel shows Healthy controls; right panel shows TLE patients. Rectangle markers represent recalled events with spoken recall time on the x-axis and film timeline on the y-axis. The five films are represented by five different colors. **B. Temporal correspondence between spoken recall and presentation order for fine film events. C. Film level recall**. Left: Recall probability per film; Right: Recall sequence error per film. Circles indicate participants with violin plots summarizing group distribution. No significant differences between HC and TLE in the number of films or film events remembered (all p > 0.05). **D. Event recall probability. Left:** Coarse events, **Right:** Fine events. No significant group differences in recall probability in either category. **E. Recall sequence errors**. TLE patients made more errors in remembering sequences of detailed film events compared to HCs (0.15±0.13 vs 0.23±0.18, p=0.027). There was a near-significant difference between TLE and HC patients in remembering the sequence of coarse events (0.10± 0.10 vs 0.19±0.18, p=0.06). **F. Sequence error by TLE lateralization**. Left: RTLE patients demonstrate more sequence recall errors of coarse events compared to HCs, although this does not reach significance (0.10±0.10 vs 0.24±0.21, p = 0.051). Right: RTLE patients demonstrate more errors in sequence memory for fine events compared to HCs (0.15± 0.13 vs 0.29± 0.21, P = 0.021); other pairwise comparisons are not significant. Additional figures are shown in **Supplementary Figures 1-2**.

**Figure 3.**
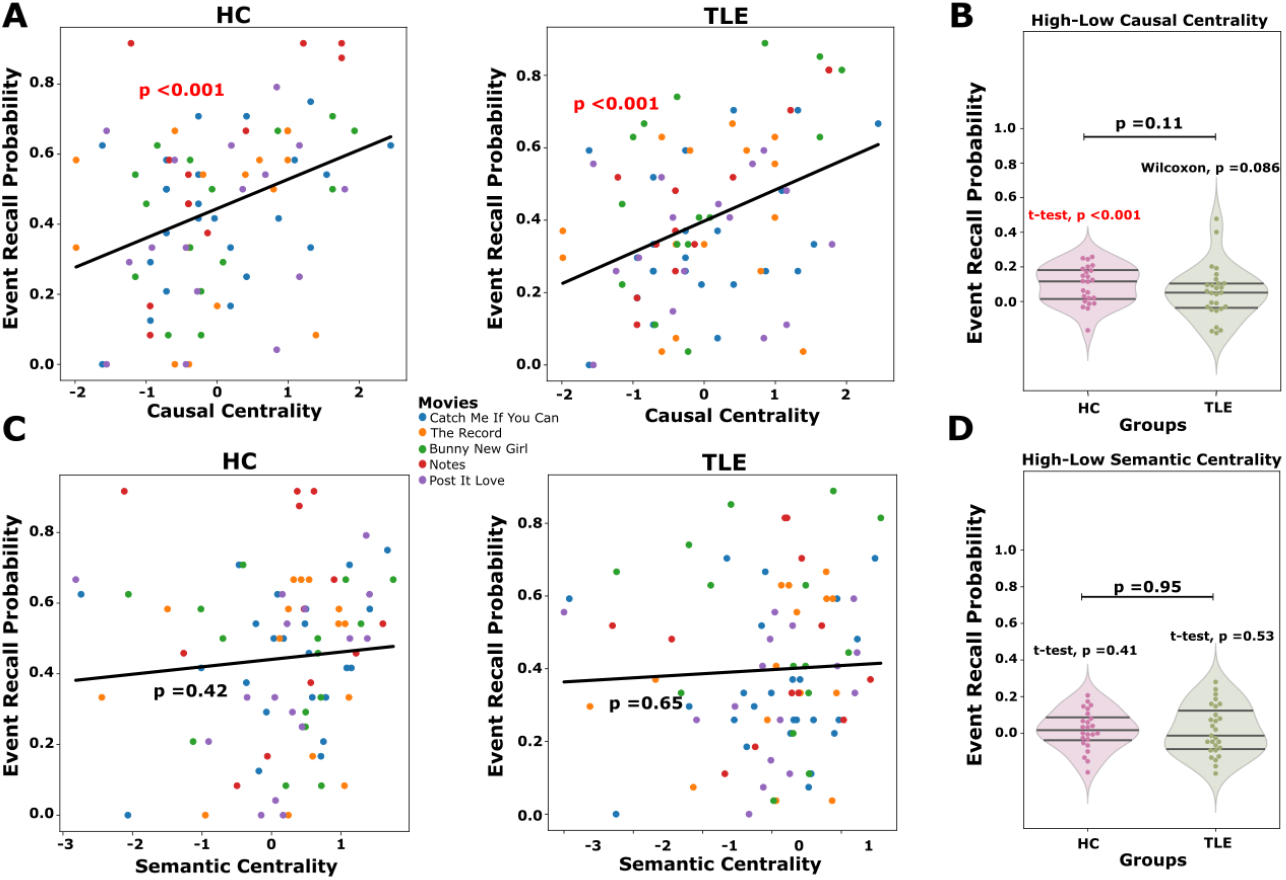
Centrality Analysis. A. Events with higher causal centrality are better remembered by HCs and TLE patients (R= 0.129, p<0.001; R=0.153, p<0.001, respectively. B. High causal centrality events are remembered better than low causal centrality events by both HCs and TLE patients. C. Higher semantic centrality does not predict recall by either HCs (p=0.42) or TLE patients (p=0.65). D. Events with high and low semantic centrality are remembered similarly by HCs and TLE patients (p=0.41 and p=0.53, respectively).

Post-hoc pairwise analysis suggests that the differences in fine event sequence recall error are driven by patients with RTLE. HCs had better sequence memory than RTLE patients (0.15 ±0.13 vs 0.29 ±0.21, p = 0.021, Hedges’ *g* = 0.85, Cliff’s δ = 0.51) (**Figure 2F**). Such differences were not found between HCs and LTLE patients or between LTLE and RTLE patients. When restricting analysis to compare only TLE patients with higher MOCA scores (≥26) to HCs. TLE patients trended toward worse sequence memory compared to HCs (HC 0.15 ±0.13 vs TLE 0.23 ±0.17, p = 0.066, Cliff’s δ = 0.327). Finally, we did not find a difference in recall probability for film F=0.0674, p=0.7056), coarse event (F=1.2337, p=0.3003), or fine event memory (F=1.6465, p=0.2034) between HC, LTLE, and RTLE patients, see **Supplementary Figure 3**.

### Recall of Film Events depends on TLE diagnosis and is independent of age, MOCA Score, or testing modality

MOCA scores did not predict sequence memory performance in HCs or TLE patients for coarse events (HC: p = 0.87; TLE: p = 0.24) (**Figure S2 A**) or fine events (HC: p = 0.51; TLE: p = 0.42) (**Figure S2 B**). TLE patients were categorized into high MOCA (scores ≥26) and low MOCA (scores <26) groups. Among TLE patients, those with high vs low MOCA scores did not differ significantly in fine event recall (0.23 ±0.11 vs 0.21 ±0.12, p = 0.51) or fine event sequence performance (0.18 ±0.15 vs 0.26 ±0.21, p = 0.31) (**Figure S2**). MOCA did not predict sequence memory performance (p = 0.459), nor did age (p = 0.640) or testing condition (in person vs. remote digital) (p = 0.637). In a beta-regression analysis, the TLE group showed significantly greater sequence error for fine events than Healthy controls (β = 0.794, p = 0.038, Supplementary Table 3). Even after controlling for MOCA delay score, age, and remote testing, the TLE group still exhibited more sequence memory errors than Healthy controls (β = 0.764, p = 0.030, Supplementary Table 4). We did not find a difference between first and last film recall in our TLE patients or HCs (p=0.629, see figure below, will be added to **Supplementary Figure 5**). We suspect that this is because TLE and (HC) patients had near ceiling performance on event recall.

### Comparison of Sequence Memory for Fine Events with Standard Neuropsychological Testing

A subset of our TLE patients had clinical neuropsychological testing performed for clinical reasons. For that group, we performed a beta regression analysis to determine whether sequence memory was correlated with standardized testing. Fine event sequence memory did not correlate with MOCA Delay (p=0.356), LMI (p=0.796), LMII (p=0.963), RAVLT Total (p=0.697), RAVLT VI (p=0.315), RAVLT VI (p=0.444), RCFT Delay (p=0.136) or MAC-E Total (p=0.442). Interestingly, sequence memory trended with VRII (p=0.071), another test traditionally believed to depend on non-dominant memory dysfunction (37,38).

### TLE Patients and HCs remember events with high causal centrality

We have previously shown in healthy individuals that events with higher centrality are better remembered (15). Events with high semantic and causal relationships to other events within a film are more memorable. As memorability is a feature of the film’s internal structure, we anticipated that TLE patients would behave no differently than HCs. We tested whether recall probability differed as a function of event centrality (High vs. Low) and participant group (Healthy vs. TLE) using beta regression with cluster-robust standard errors. Recall probability was significantly higher for high-centrality events than for low-centrality events, indicating a robust main effect of centrality (χ^2^(1) = 14.52, p < .001). There was no main effect of group (χ^2^(1) < 0.01, p = .98), and no evidence of a centrality × group interaction (χ^2^(1) = 0.11, p = .74). In contrast, recall probability did not differ between high- and low-semantic centrality events, indicating no main effect of semantic centrality (χ^2^(1) < 0.01, p = .998). There was no main effect of group (χ^2^(1) = 0.15, p = .70) and no evidence of a semantic centrality × group interaction (χ^2^(1) = 0.08, p = .78). Note that causal centrality was defined only for coarse events and not fine (see Methods).

## DISCUSSION

Episodic memory is commonly described as “mental time travel,” wherein personally experienced events bound to a spatial and temporal context are revisited (1). Temporal lobe epilepsy (TLE) often leads to limbic system dysfunction, including the hippocampus and partner regions, and thus impairs episodic memory (39). Results from fMRI studies showing reorganization of memory functions demonstrate changes in temporal and extratemporal regions (40). Traditional neuropsychological tools that rely on memorizing unstructured information, such as word and number lists, often fail to capture the complexity of real-life episodic memory and depend heavily on recruitment of extra-temporal brain regions including frontal executive systems. To address this, our naturalistic film paradigm mimics real-life experiences and remembering, enhancing ecological validity and providing a more challenging and multi-faceted assessment of episodic memory.

Our results show that the naturalistic film task elicits a profile of spontaneous recall in TLE patients that resembles HC recall of film events; TLE patients remembered a similar percentage of films and events compared to HCs, and events with high causal centrality were better remembered by both groups. However, TLE patients exhibited more sequence errors during recall. Importantly, these sequence recall deficits were not explained or moderated by overall cognitive status, indicating a possible disruption in temporal organization or episodic structuring not captured by broad cognitive screening tools such as the MOCA. Similarly, our findings were independent of performance on standard verbal and visual memory tasks, and distinguished between left and right TLE, suggesting that our film recall task may detect a unique and clinically relevant aspect of episodic memory.

At the group level, our TLE cohort had similar recall performance for film event details compared to HCs when tested immediately after film presentation. These findings may be due to milder memory impairment in our patients compared to previously described cohorts, as our threshold for inclusion was MOCA ≥ 22. Patients’ relatively strong event recall on the film task may also reflect superior ability for remembering structured compared to unstructured verbal information (i.e. stories versus word lists) (41). Recall of unstructured information may depend more on frontal-executive deficits rather than episodic memory deficits per se (4). Furthermore, we asked subjects to recall films immediately after viewing, contributing to a near ceiling effect at film- and event-level recall.

While event-level recall was comparable to HCs, TLE patients demonstrated poorer recall for the temporal order of finely segmented events, and a near significant difference for coarse events. which may reflect a more subtle decline in episodic memory function or a distinct behavior altogether. While the “what” and “where” aspects of episodic memory have been well-studied, the temporal features of recall, particularly sequence memory, remain less well-characterized inpatient groups such as temporal lobe epilepsy. Notably, sequence memory has been explored in the context of Alzheimer’s disease, with tasks like picture sequence memory and logical memory (i.e., Craft Story recall) sensitive to early pathological change (42). Patients with severe hippocampal damage have been demonstrated to have poorer recall of specific event details as well as impaired memory for the order of events (43). The degree of immediate memory loss for event details in patients with hippocampal damage has been demonstrated to resemble healthy forgetting at one month after the event (44).

Our finding that sequence memory was slightly worse in RTLE patients compared to LTLE patients is intriguing and merits follow-up investigation. While thought to reflect the “material specific” function of left versus right temporal lobe dysfunction, the RCFT has been criticized for its inability to distinguish between L and RTLE patients (45). This may be due to the confounding influence of verbal learning and executive strategies commonly employed to draw and recall the figure. We note that sequence performance trended with RCFT delayed recall scores (p=0.07), but not MOCA scores, or verbal and visual memory scores in the subset of our patients who had neuropsychological testing. This unique finding needs to be replicated in larger patient cohorts, but could be useful in clinical settings to detect non-dominant temporal lobe dysfunction.

We note that an event’s causal centrality strongly predicted recall performance in both groups, while semantic centrality had a weaker or inconsistent effect. This finding is consistent with previous research demonstrating that causal connections are particularly critical for narrative memory in healthy patients (46); for example, a recent study showed that individuals rely on a causal strategy for recalling narratives, even when explicitly instructed to avoid doing so (19). In another study, it was found that the impact of semantic centrality on narrative memory could be modified by experimental manipulation (in this case, the sense of agency), while the impact of causal centrality on memory was unaffected (47). Causal processing may serve as a conserved strategy for organizing episodic narratives, and this could be reflected in the resilience of causal connections even in memory-impaired populations. In contrast, semantic knowledge is influenced by age, education, and task-related factors, which may explain why semantic centrality was a variable predictor of later memory.

### Prior work on the neural underpinnings of sequence memory

From a cognitive neuroscience perspective, the sequence memory impairments observed in TLE patients may stem from disruptions in scene construction and/or retrieval of temporal context. Scene construction, or the act of re-creating coherent mental representations of events anchored in time and space, depends on interactions between hippocampus and ventromedial prefrontal cortex (48). This ability to re-construct prior events is compromised in patients with medial temporal lobe damage (49). While the TLE patients in our study did not demonstrate a deficit in basic event recall, they did have impaired recall of event order which requires a greater memory demand. Sparse memory for an event could be sufficient to be called “remembered.” However, more details — particularly details that link events—are needed to recover temporal order. Thus, deficits in scene construction may have contributed to TLE patients’ weaker scaffolding to accurately recall the temporal order of events. Furthermore, studies have shown that the MTL plays a critical role in retrieving temporal context, with hippocampal and MTL cortical activity patterns encoding the temporal position of objects in learned sequences (50). In line with this idea, temporal context is impaired in patients with MTL lesions (51). Sequence memory may arise from disruptions in hippocampal and prefrontal function, or communication between these regions, degrading the temporal organization of episodic content. Because temporal order memory appeared to be worse in RTLE compared to LTLE, we hypothesize that the difference in our subgroups was due to a hippocampal deficit.

There are several potential neurophysiological mechanisms to support sequence memory. Hippocampal CA3 association across the long-axis of the hippocampus appears to be critical for maintaining temporal order in mice, with selective silencing disrupting acquisition order, but not to object or location-related information (52). Neurons in monkey entorhinal cortex exhibit increased firing rate in response to visual stimuli, with relaxation occurring at different time intervals, facilitating temporal discrimination. Hippocampal theta-gamma phase amplitude coupling (PAC) has been established as a core mechanism of encoding place cell order during spatial navigation in rodents (53), While there is some evidence that these mechanisms may be at play in humans (54), these findings require further investigation, including how these cardinal oscillations and subfield-specific functions may be disrupted in neurological conditions such as TLE and Alzheimer’s Disease.

### Limitations and Future Research Directions

We acknowledge the limitations of our exploratory study. One weakness is the limited validation against standard neuropsychological testing. Thus, we cannot claim that film task stimuli and associated recall are more sensitive or specific than traditional neuropsychological tests. Future studies could include a larger battery of standard neuropsychological tests for both healthy and TLE patients, larger numbers of patients, and inclusion of other focal and generalized epilepsy populations, to understand how “cognitive phenotypes” are differently expressed when recalling films. Another limitation of our study includes limited sampling of subjective memory impairment, to determine whether early decline can be detected by a film task. Furthermore, while films require memory for a greater amount of information compared to standard lists or paragraphs, both HCs and patients performed near ceiling on film and event recall. A task with more films, longer films, or delayed recall would likely show a greater range of performance, potentially at immediate recall or after a brief distraction.

While we did not include delayed recall of >30 minutes into our task, we would expect that delayed recall would demonstrate larger effect sizes. Some traditional neuropsychological tests can demonstrate only memory deficits in TLE patients with subjective memory complaints after a delay of days to weeks (23). We note that accelerated forgetting is a widely described phenomenon in TLE patients (55–61); and that future studies might explore how event and sequence memory decay at different rates. Replication of our findings of poorer sequence memory in RTLE compared to LTLE patients in larger patient cohorts is necessary. Future studies with both immediate and delayed recall testing would clarify how different facets of episodic memory (e.g. who, what, where, when) decay at different rates.

### Clinical implications

The immediate clinical implication is that naturalistic episodic memory tasks, such as in film recall could be deployed as a complementary means of episodic memory testing. Film recall requires subjects to associate the who, what, when, where, and why of an event, similar to everyday recall behavior. Some commonly administered neuropsychological tests, such as word or story recall tasks (RAVLT, CVLT, WMS LM, & Craft Memory Test), are used to assess memory for unstructured and structured verbal information. While the number of words or units of information remembered is scored in neuropsychological assessment, there is inconsistent use of sequence information.

A practical first step may be to modify or extend analysis of existing tests – such as focusing on serial order of word recall in list-learning tests or analyzing spontaneous recall of narrative details in story recall tasks. Recent efforts, such as the NIH Cognitive Toolbox Picture Sequence Memory Task, have aimed to systematically evaluate sequence memory. Already, impaired memory for early story events has already demonstrated mild cognitive impairment and amyloid burden in cognitively unimpaired patients (61). EpiReal is another novel task designed to capture more naturalistic elements of autobiographical recall, presents 8 mini events to subjects (e.g. “hand the examiner a green binder located on the chair”) (23). Besides probing memory for the content of the event, the task requires memory for spatial and temporal context. EpiReal has been demonstrated to be sensitive to accelerated long-term forgetting in TLE patients, with some evidence that it could be more sensitive than standard list learning tasks. Our film task resembles EpiReal in its ability to test multi-faceted events (although not personally experienced events), with the additional dimension of temporal sequence order.

## CONCLUSION

In summary, a naturalistic film recall task demonstrates that TLE patients recall a similar percentage of films and film events compared to healthy controls, yet make more sequence errors in recalling detailed events. These findings suggest that naturalistic task stimuli, such as film recall, can discover unique impairments in episodic memory that are typically unmeasured in clinical settings. Future studies should replicate and validate our findings within larger cohorts of TLE patients and compare performance on a film series task to standard neuropsychological testing performance.

## ACKNOWLEDGEMENTS

This work was supported by NIH grants R01NS127954 (AL) and K23NS104252 (AL). Herui Zhang and Forouzan Farahani contributed equally to the project. Janice Chen and Anli Liu contributed equally to the supervision of this project.

## CONFLICT OF INTEREST

None of the authors has any conflict of interest to disclose.

## Notes

### Competing Interest Statement

The authors have declared no competing interest.

