## Supplementary for "Film Recall Reveals Intact Event Memory but Impaired Sequence Memory in Temporal Lobe Epilepsy Patients"

**Instructions: Causality labeling**

You will watch up to 5 short movies, one at a time. For each movie, we would like you to identify **causal relationships** between events.

We want to know how causal relationships between the events within a movie affect the brain responses and memory performance for the events.

In the folder for each movie, you will also find a spreadsheet with descriptions for the movie, segmented into “events” and their finer-grained “sub-events”. Your job is to identify and make a list of event pairs that are causally related to each other within each movie.

Download the ‘causality.xlsx’ and open the file on your local machine. For each row, enter the event ID numbers of the event pair with a cause-effect relationship as below. **ONLY enter event ID, NOT sub-event ID.**

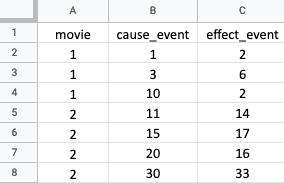

Most events consist of more than one sub-events. Consider two events as causally related if any sub-event of an event is the cause of any (at least one) sub-event of the other event. An event pair should always consist of two *different* events. That is, ignore the causal relationship between sub-events of the same event.

Note that the movie event order does not always follow the actual chronological order of events occurred in the story. In other words, higher-numbered events can cause lower-numbered events.

How can we decide whether two events are causally related or not? In an extremely broad sense, one might say that any event that happened before a target event could be at least partially responsible for the event to happen (e.g., you were born because there was Big Bang), but this wouldn’t give us very useful information. So we want to identify only those event pairs that are more strongly related, and you will need to use your own best judgment to decide whether the causal relationship is strong enough. For example, if we have a movie like below,

Event 1: Jane orders a crab cake at a restaurant.

Event 2: Jane finds a dead fly in her crab cake.

Event 3: Jane complains to the manager of the restaurant.

You may say that there is a causal relationship between Event 2 and Event 3, but not between Event 1 and Event 3. We don’t really have strict rules or criteria, so it is up to your subjective judgment. But please try to keep your criteria as consistent as possible.

Complete one movie at a time. That is, open one folder and watch the movie inside; then review the Segmentation sheet; then fill out the downloaded “causality.xlsx” for that movie.

When you are finished with one movie, proceed to the next one. We will check in with you periodically. Take your time—carefulness is much more important than speed.

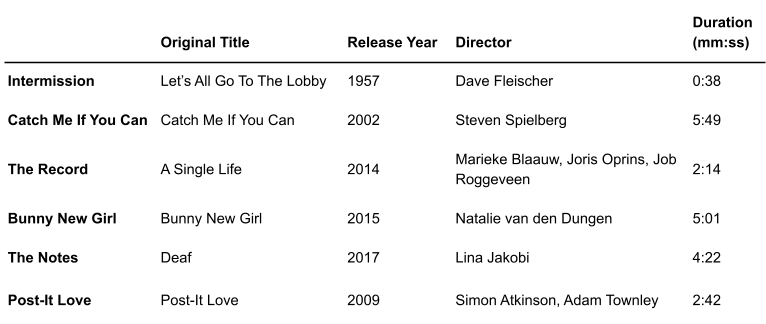

**Supplementary Table 1. Film Description**

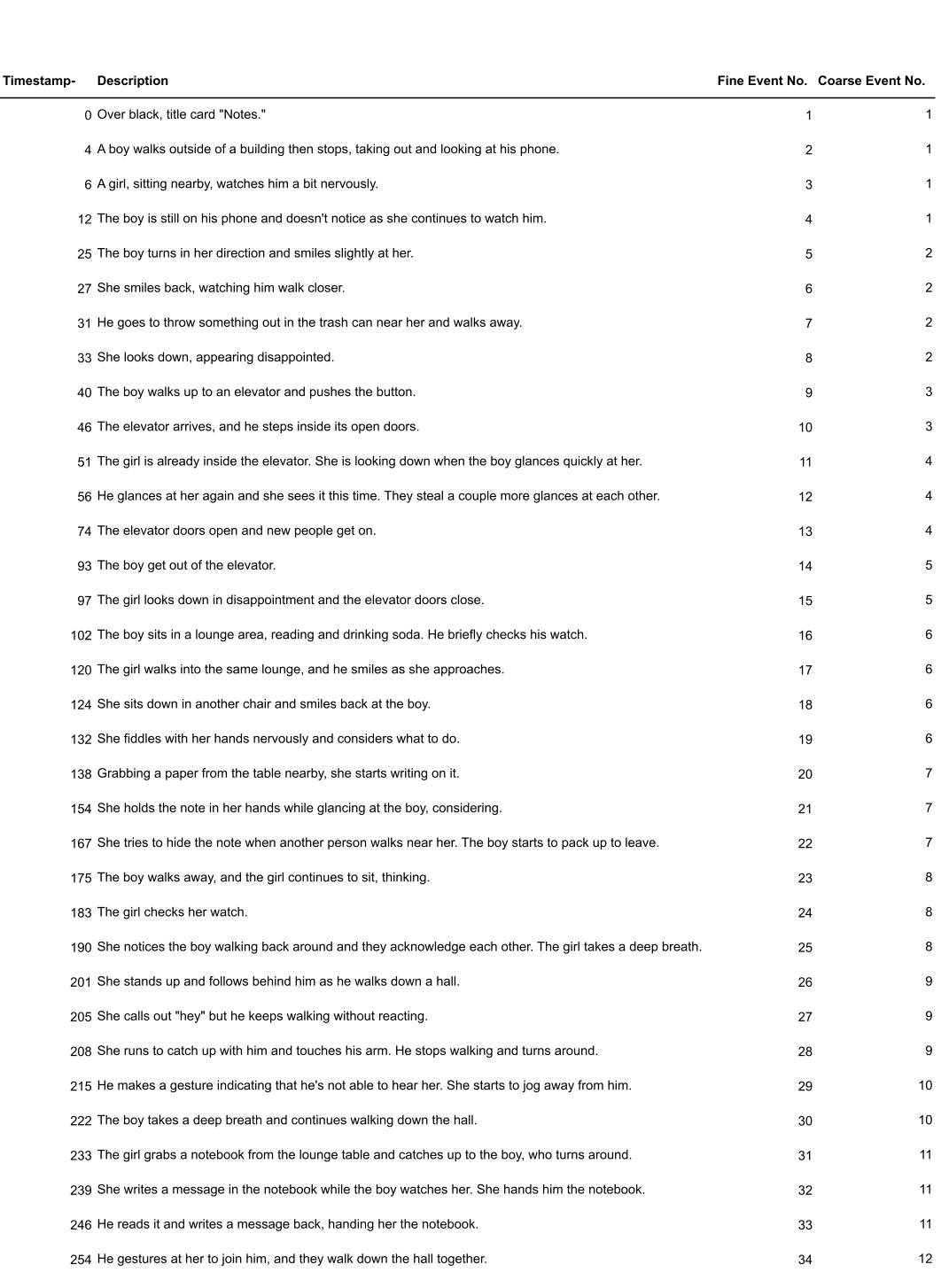
**Supplementary Table 2. Example fine event boundaries for “The Notes.”**

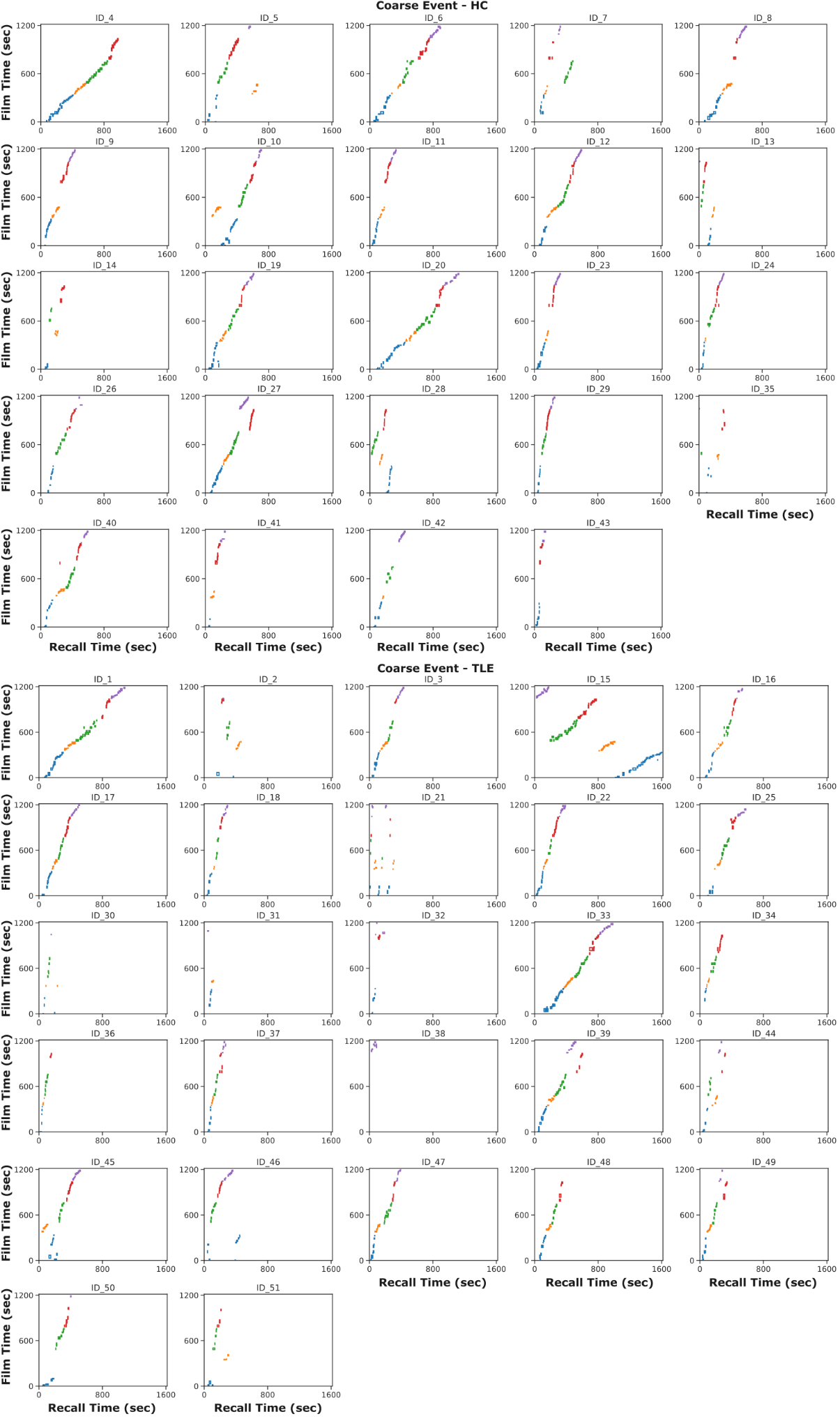

**Supplementary Figure 1. Healthy Control and Temporal Lobe Epilepsy Patients Recall of Coarse Film Event Segments.**

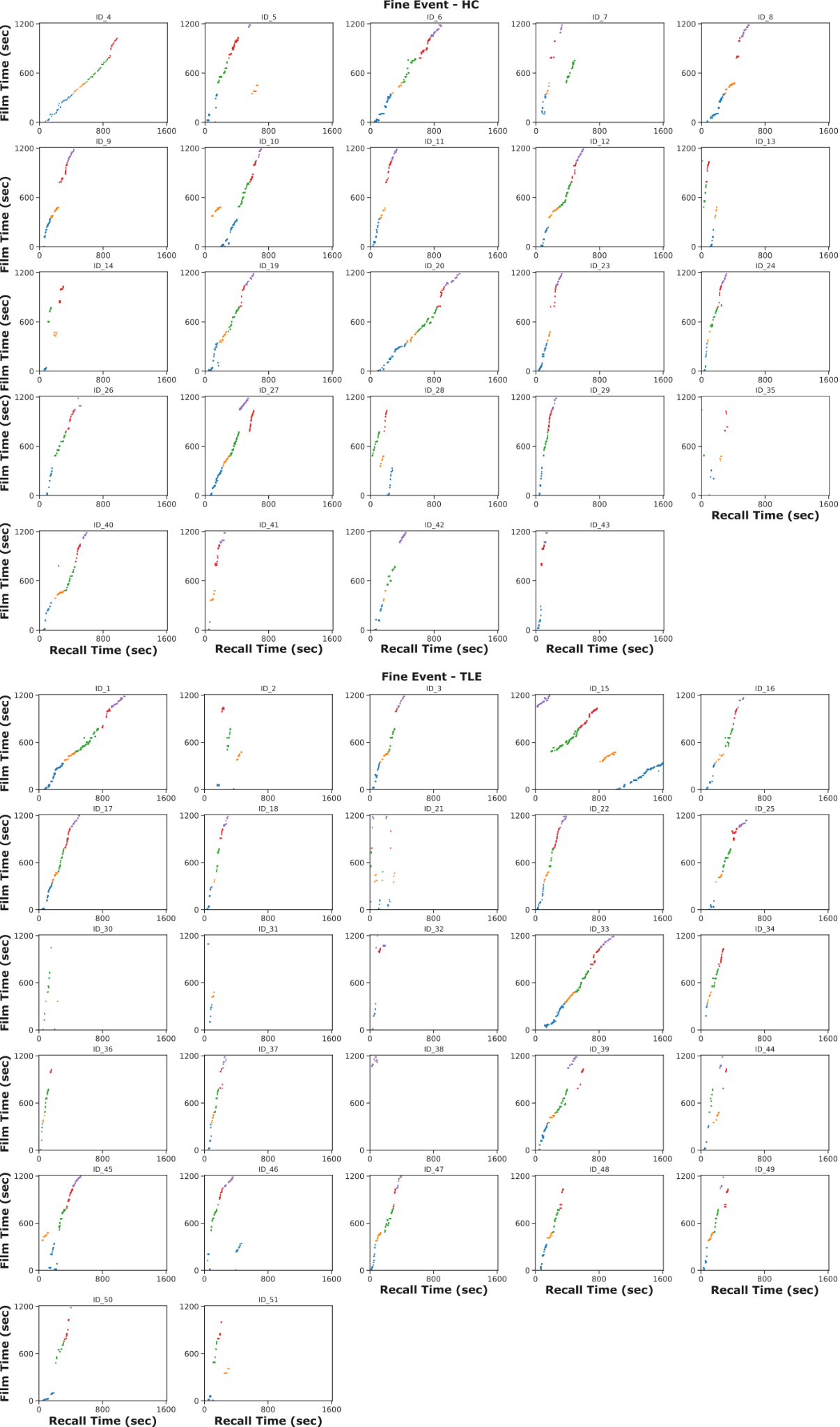

**Supplementary Figure 2. Healthy Control and Temporal Lobe Epilepsy Patients Recall of Fine Film Event Segments.**

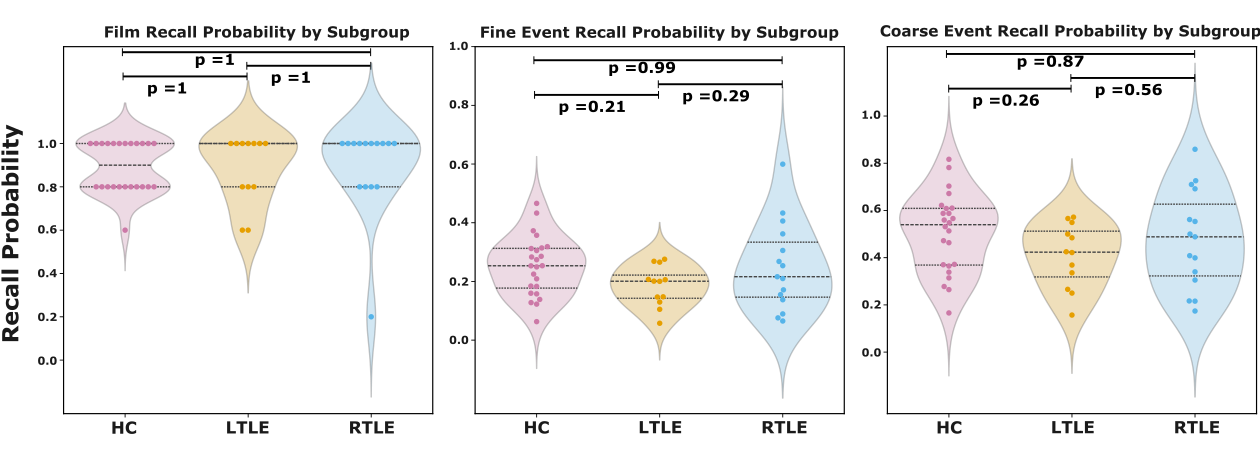

**Supplementary Figure 3. Recall Probability by TLE lateralization.**

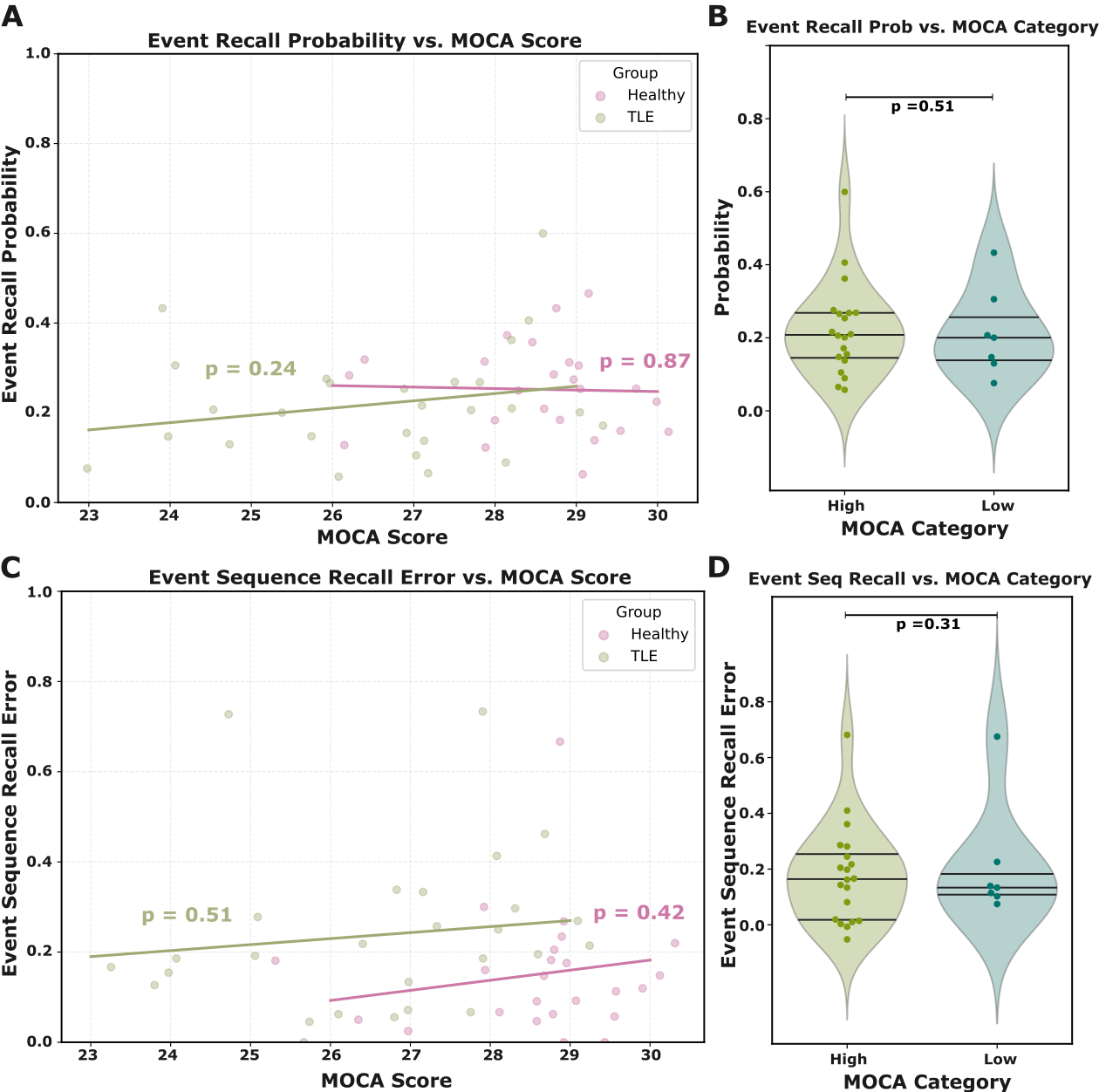

**Supplementary Figure 4. A: Recall probability vs. MOCA.** Each point is a participant. X-positions include small horizontal jitter to reduce overlap. Lines show separate best-fit trends for each group. **B: Recall probability by the MOCA group.** Violin plots show the distribution of recall probability for Low vs High MOCA participants (TLE only). **C: Recall Sequence Error vs. MOCA.** Each point is a participant. X-positions include small horizontal jitter to reduce overlap. Lines show separate best-fit trends for each group. **D: Recall sequence error by the MOCA group.** Violin plots show the distribution of recall sequence error for Low vs High MOCA participants (TLE only).

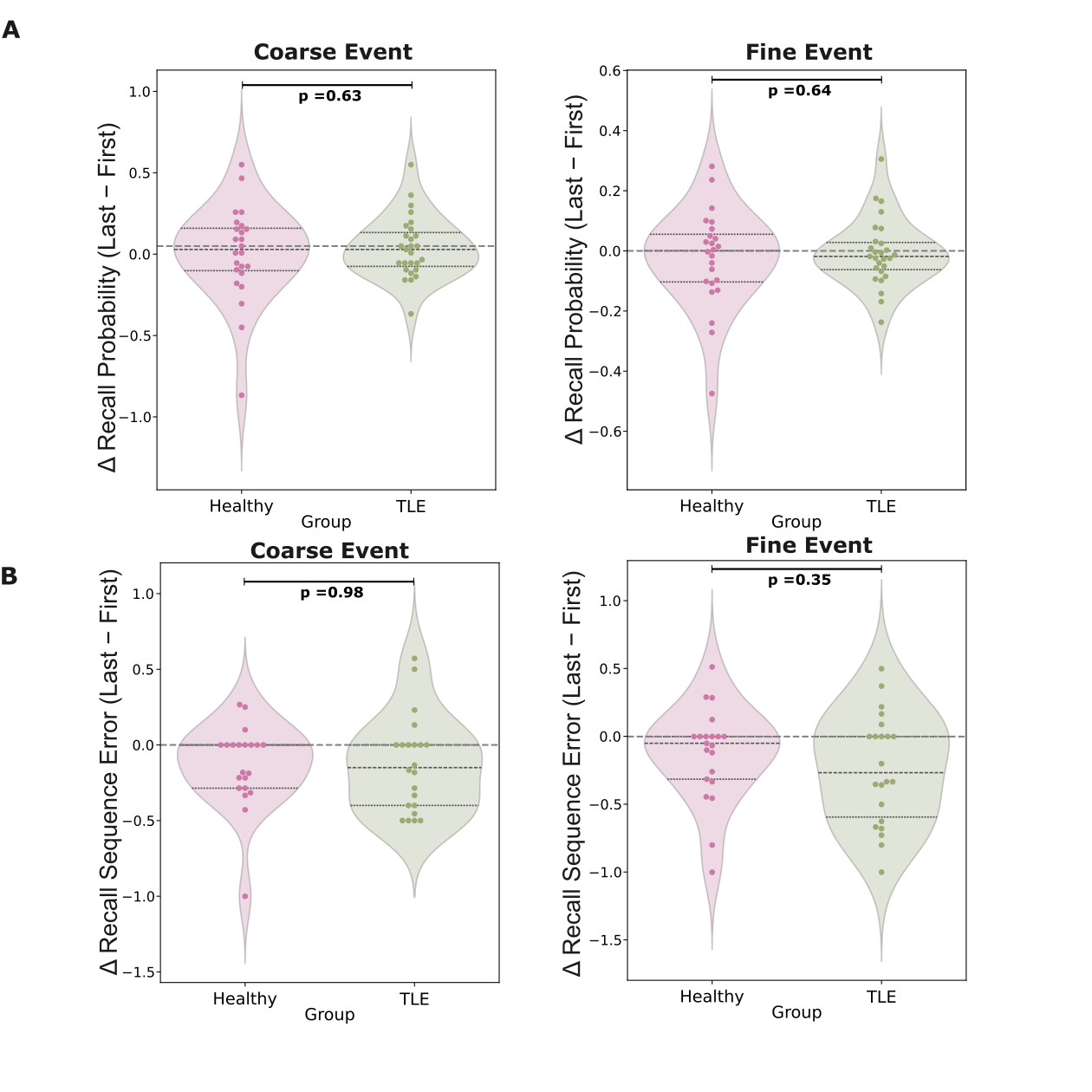

**Supplementary Figure 5. A: Change in Recall Probability (Last − First):** Violin plots show the distribution of Last − First recall for HC and TLE. Positive values indicate higher recall for the last film; negative values indicate a decline. Individual points represent participants. No significant difference between HC and TLE. **B: Change in Recall Sequence Error (Last − First):** Violin plots show the distribution of Last − First recall for HC and TLE. Positive values indicate higher recall for the last film; negative values indicate a decline. Individual points represent participants.

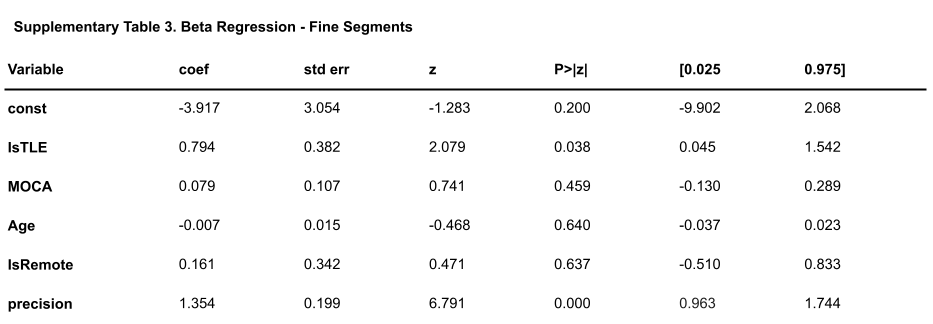

**Supplementary Table 3. β-regression of recall sequence error.** Recall sequence error was modeled as a function of TLE status, platform (in-person vs. remote), age, and MOCA.

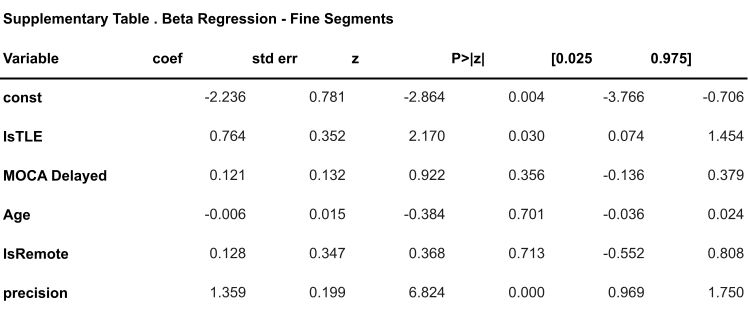

**Supplementary Table 4.β-regression of recall sequence error.** Recall sequence error was modeled as a function of TLE status, platform (in-person vs. remote), age, and MOCA delay.

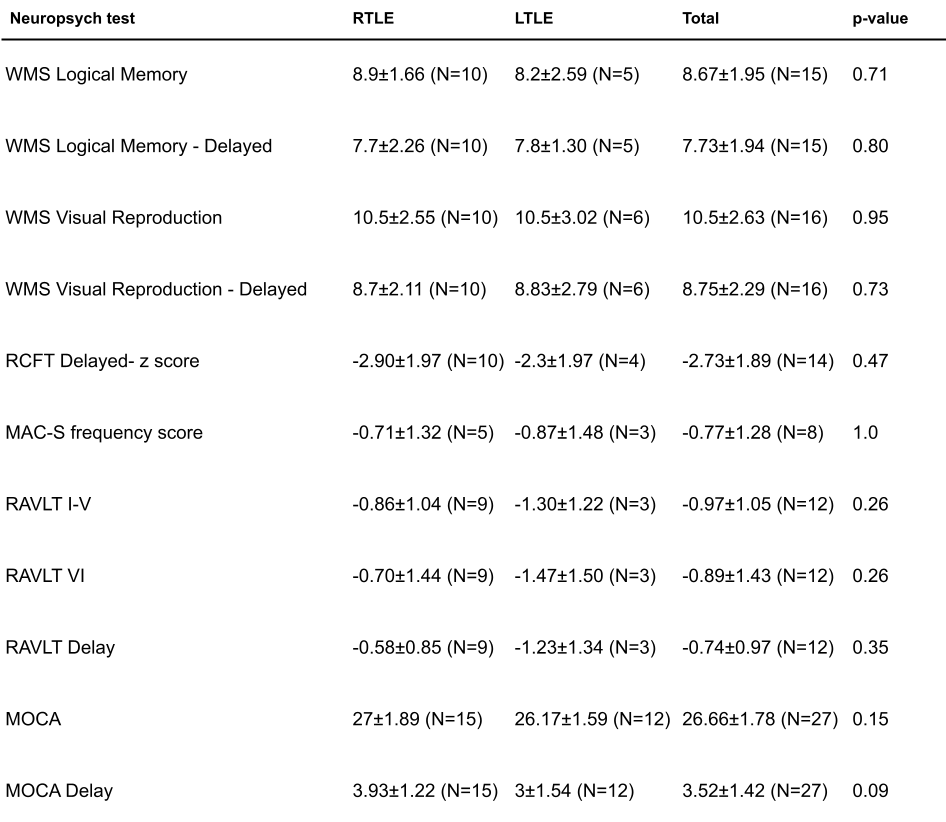

**Supplementary Table 5. Summary of neuropsychological test scores in RTLE vs. LTLE. Values are mean ± SD (N) for each measure.**

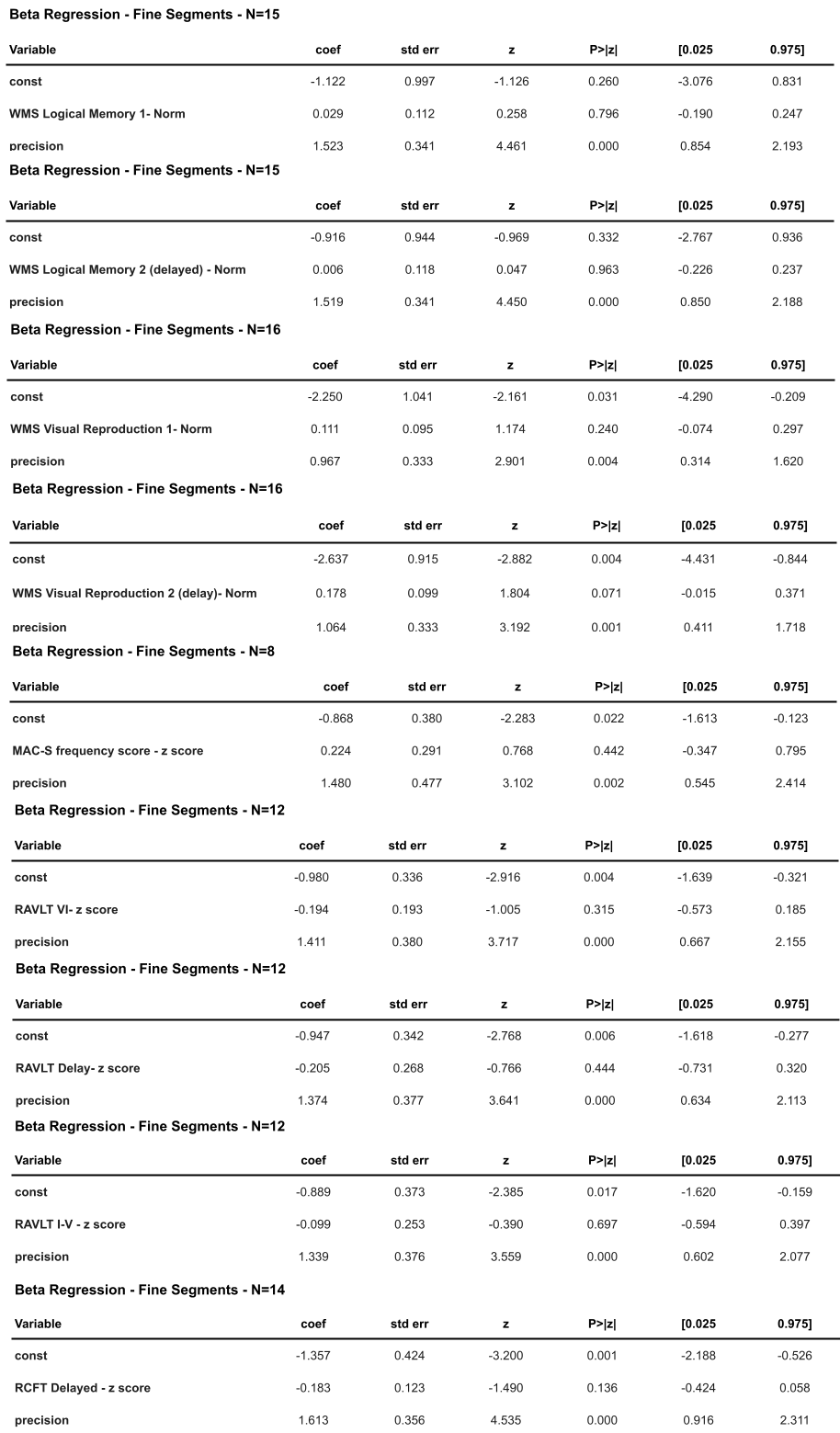

**Supplementary Table 6. Effects of neuropsychological test performance on recall sequence error (β-regression).** Entries are coefficient (coef), standard error (std err), p, and 95% CI. Negative coefficients indicate that higher test scores predict fewer sequence errors.
